# Increased pathogenicity and transmission of SARS-CoV-2 Omicron XBB.1.9 sublineage, including HK.3 and EG.5.1

**DOI:** 10.1101/2024.06.10.598324

**Authors:** Qiushi Jin, Ruixue Liu, Wenqi Wang, Jichen Xie, Tiecheng Wang, Haiyang Xiang, Xianzhu Xia, Jianmin Li, Xuefeng Wang, Yuwei Gao

## Abstract

With the SARS-CoV-2 Omicron XBB.1.9 sublineage circulating worldwide, two XBB.1.9 variants, EG.5.1 and HK.3 spread rapidly and became dominant from middle 2023. However, the spike features, pathogenicity, and transmissibility of HK.3 are largely unknown. Here, we performed multiscale investigations to reveal the virological features of XBB.1.9 subvariants, especially the newly emerging HK.3. HK.3 revealed high replication efficiency in vitro. The HK.3 spike exhibited enhanced processing, although its infectivity, fusogenicity, and hACE2 binding affinity were comparable to those of the EG.5 and XBB.1 spikes. All XBB.1.9.1, EG.5.1 and HK.3 strains demonstrated efficient transmission in hamsters, although XBB.1.9.1 exhibited stronger fitness in the upper airways. HK.3 and EG.5.1 exhibited greater pathogenicity than XBB.1.9.1 and BA.2 in H11-K18-hACE2 hamsters. Our studies provide insight into the newly emerging pathogens HK.3 and EG.5.1.

**Importance:** In animal models, the ongoing attenuated pathogenicity and poor transmission of Omicron subvariants seems to reach a consensus. However, our results revealed that Omicron XBB.1.9 subvariants, including one of the key variants of interest, EG.5 with its another key subvariant HK.3, universally exhibited both increased pathogenicity and highly transmission. This study reemphasized the importance of surveillance in characteristics of epidemic Omicron subvariants.

## Introduction

The coronavirus disease 2019 (COVID-19) pandemic still lingers globally. One of the greatest challenges during the COVID-19 pandemic was the speed at which the causative agent severe acute respiratory syndrome coronavirus 2 (SARS-CoV-2) mutated. The Omicron BA.1 variant emerged in November 2021 and was characterized by more than 30 new mutations in the spike alone, and subsequent Omicron sublineages have continued to accumulate additional mutations (1). Identified in September 2022, the XBB lineage originated from a recombination of two BA.2-derived variants (BJ.1 and BM.1.1.1), mainly included XBB.1.5, XBB.1.9, XBB.1.16 and XBB.2.3, and progressively replaced most of the previous Omicron strains. These variants exhibit notable changes in virological characteristics, including increased transmissibility and obvious immune evasion (2–5).

EG.5.1, a subvariant representing most EG.5 strains (XBB.1.9.2.5), quickly spread in several areas of the world and replaced the previous XBB.1.5, XBB.1.9, and XBB.1.16 variants, which also evade most XBB.1.5-neutralizing antibodies (6). EG.5.1 has further evolved into a descendant lineage bearing the spike L455F mutation and has been named HK.3 (XBB.1.9.2.5.1.1.3), which showed increased transmission (7).

In this work, we investigated the in vitro and in vivo virological characteristics of isolates of XBB.1.9 subvariants. The in vitro replicative kinetics of XBB.1.9.1, EG.5.1 and HK.3 were compared with those of the former epidemic strain BA.2. The pathogenicity of XBB.1.9.1, EG.5.1 and HK.3 in wild-type and K18-hACE2 rodents was determined. We also demonstrated the airborne transmissibility of XBB.1.9.1, EG.5.1 and HK.3 in hamsters. Moreover, as the spike protein universally impacts the infection and pathogenicity of SARS-CoV-2 via its functional characteristics (8–11) changes in spike features were also studied.

## Results

### Prevalence and mutations of XBB.1.9 subvariants

Compared to XBB.1, XBB.1.9 bears the amino acid mutations G1819S and T4175I in ORF1a (Fig. 1A), with two diverging sublineages, both bearing the S486P mutation in the spike protein. One cluster was XBB.1.9.1 with the spike S486P and the nonsense C11956T mutations. The other cluster was XBB.1.9.2, which has the spike S486P mutation and the nonsense T23018C and A27507C mutations. The geographic distribution of XBB.1.9.1 and XBB.1.9.2 mainly includes Indonesia, England, the United States, and China. As a subvariant of XBB.1.9.2, EG.5.1 bears the spike Q52H/F456L double mutations, and its subvariant HK.3 contains a further L455F mutation in the spike protein. XBB.1.9.1 was one of the fastest growing variants in several areas of the world, including China, the United States, Europe, Korea and Singapore, starting in early January 2023; however, this variant was quickly replaced by XBB.1.9.2 within 2 to 3 months (Fig. 1B). Since May 2023, the prevalence of EG.5.1 and HK.3 has continued to increase in China after their emergence, accounting for up to 50% and 40% of new infections, respectively. Moreover, from July 2023 to February 2024, the relative prevalence of HK.3 reached 8%, 7%, and 52%, respectively, in the United States, Europe, and South Korea. XBB.1.9 variants exhibit notably increased transmissibility, and their remarkable immune evasion also triggered concern (7, 12, 13). Thus, we subsequently investigated the virological characteristics of XBB.1.9.1, EG.5.1 and HK.3.

**Fig. 1.**
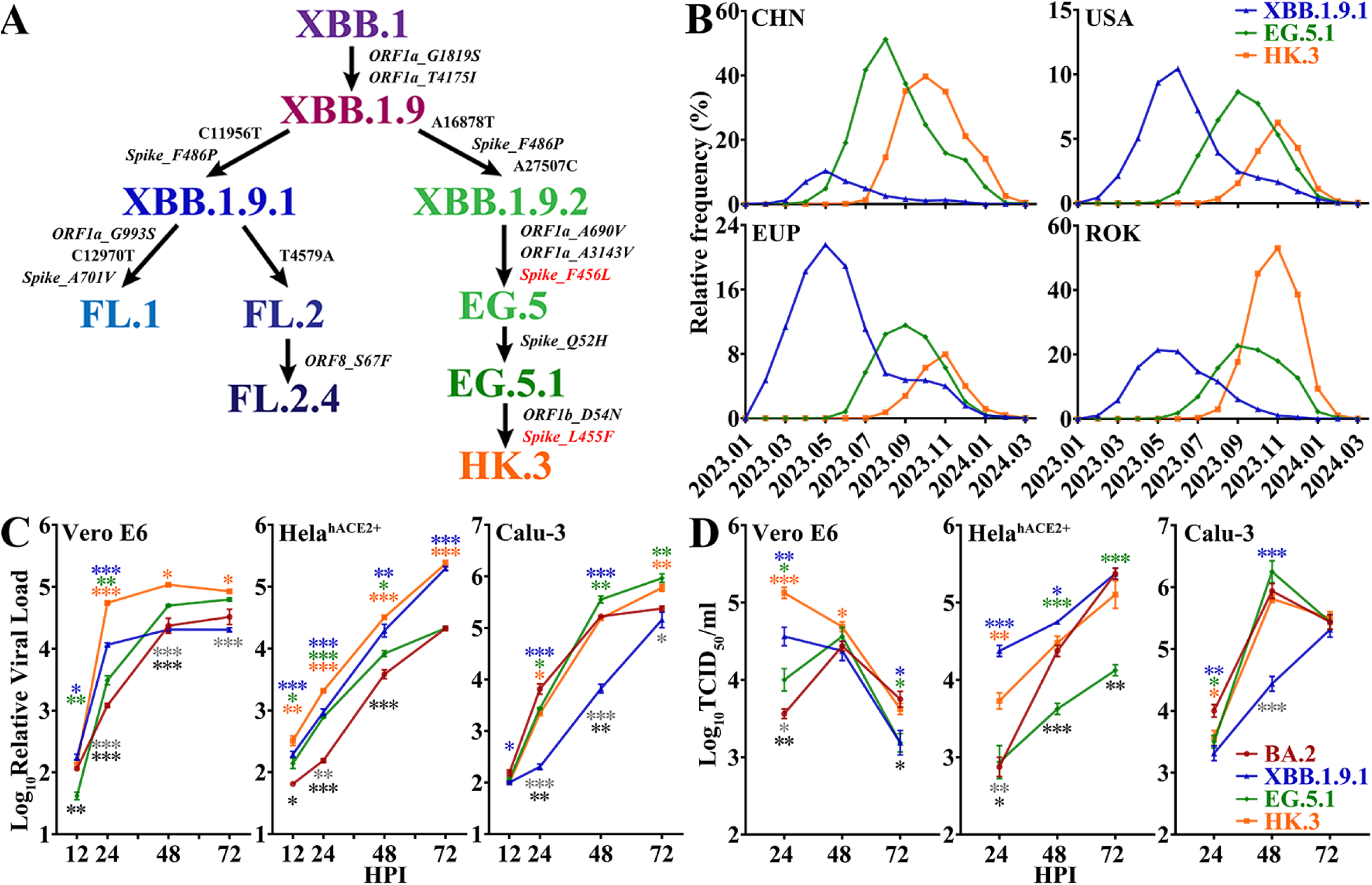
Evolution, prevalence and replicative kinetics of XBB.1.9 subvariants. A. Evolutionary origins of the XBB.1.9 sublineages, including XBB.1.9.1, EG.5.1 and HK.3. Synonymous mutations in nucleotides and amino acid mutations are shown in bold and bold-italic font, respectively. B. Prevalence of XBB.1.9.1 (blue), EG.5.1 (green) and HK.3 (orange) in China (CHN), the United States (USA), Europe (EUP) and Korea (ROK) for fourteen months from January 2023 (2023.01) to March 2024 (2024.03). C-D. Replicative kinetics of BA.2 (dark red), XBB.1.9.1 (blue), EG.5.1 (green) and HK.3 (orange) in terms of viral load (C) and viral titer (D) in Vero E6, HeLahACE2+ and CaLu-3 cells. The significance of the differences in replication between BA.2 and XBB.1.9.1, EG.5.1 or HK.3 is indicated above the lines by the asterisks in colors corresponding to the individual viruses. The significance of the differences in replication between HK.3 and XBB.1.9.1 or EG.5 is indicated by gray or black asterisks below the lines.

### In vitro replication of Omicron XBB.1.9 subvariants

The replication kinetics of BA.2, XBB.1.9.1 and the subvariants EG.5.1 and HK.3 were compared by determining viral loads and viral titers from 12 to 72 hours post infection (HPI). HK.3 replicated faster and more efficiently than BA.2, EG.5.1 and XBB.1.9.1 in Vero E6 cells (Figs. 1C and 1D). However, compared to HeLa^hACE2+^ and CaLu-3 cells, Vero E6 cells poorly supported reproduction of all the accessed Omicron strains, resulting in titers lower than 1×10^5^ TCID_50_/ml. In TMPRSS2-negative HeLa^hACE2+^ cells, HK.3 also exhibited increased replication compared with the parental EG.5.1 strain. Both EG.5.1 and HK.3 propagated well in hACE2 and TMPRSS2 double-positive CaLu-3 cells, and their replication kinetics were significantly greater than those of XBB.1.9, especially at 24 and 48 HPI, indicating that the TMPRSS2 usage in EG.5.1 and HK.3 was better than that of the parental XBB.1.9 strain. Thus, HK.3 exhibited universal replication fitness compared with its parental strains. Furthermore, few replication discrepancies between the two HK.3 isolates (CC11 and CC12) were found in the three cell lines (Fig. A1). Thus, only HK.3-CC11 was used for further studies.

### Characteristics of XBB.1.9 spike mutations

The characteristics of spike proteins are regarded as key factors in in vitro replication, pathogenicity and transmission (8, 14, 15). To investigate the mutation-induced characteristics of the XBB.1.9 spike protein, we first determined spike-mediated infectivity by pseudovirus-based infection. F486P in XBB.1.9 resulted in significantly greater infectivity than that of XBB.1 with F486S in HeLa^hACE2+^ cells (Fig. 2A). Although the further variants, EG.5.1 and HK.3, retained high infectivity, there were no significant differences between these strains and XBB.1-F486P, similar to the findings of a recent study (7). The XBB.1-F486P, EG.5 and HK.3 spikes also resulted in greater infectivity than those of D614G and BA.2. The formation of syncytia is regarded as a hallmark of SARS-CoV-2-induced pathogenesis in the lungs (16, 17) and is caused by spike-ACE2 interactions on the cell surface, referred to as spike-mediated cell‒cell fusion. At 6 hours post-cell contact, despite the large intragroup differences in both XBB.1-F486P and EG.5.1, the XBB.1-F486S and HK.3 spikes demonstrated significantly greater fusogenicity than that of BA.2 (Fig. 2B). The differences in the fusogenicity of the XBB.1-F486P, EG.5.1 and HK.3 spikes were not significant, and their expressions induced smaller syncytia than D614G and larger syncytia than BA.2 (Fig. 2C).

**Fig. 2.**
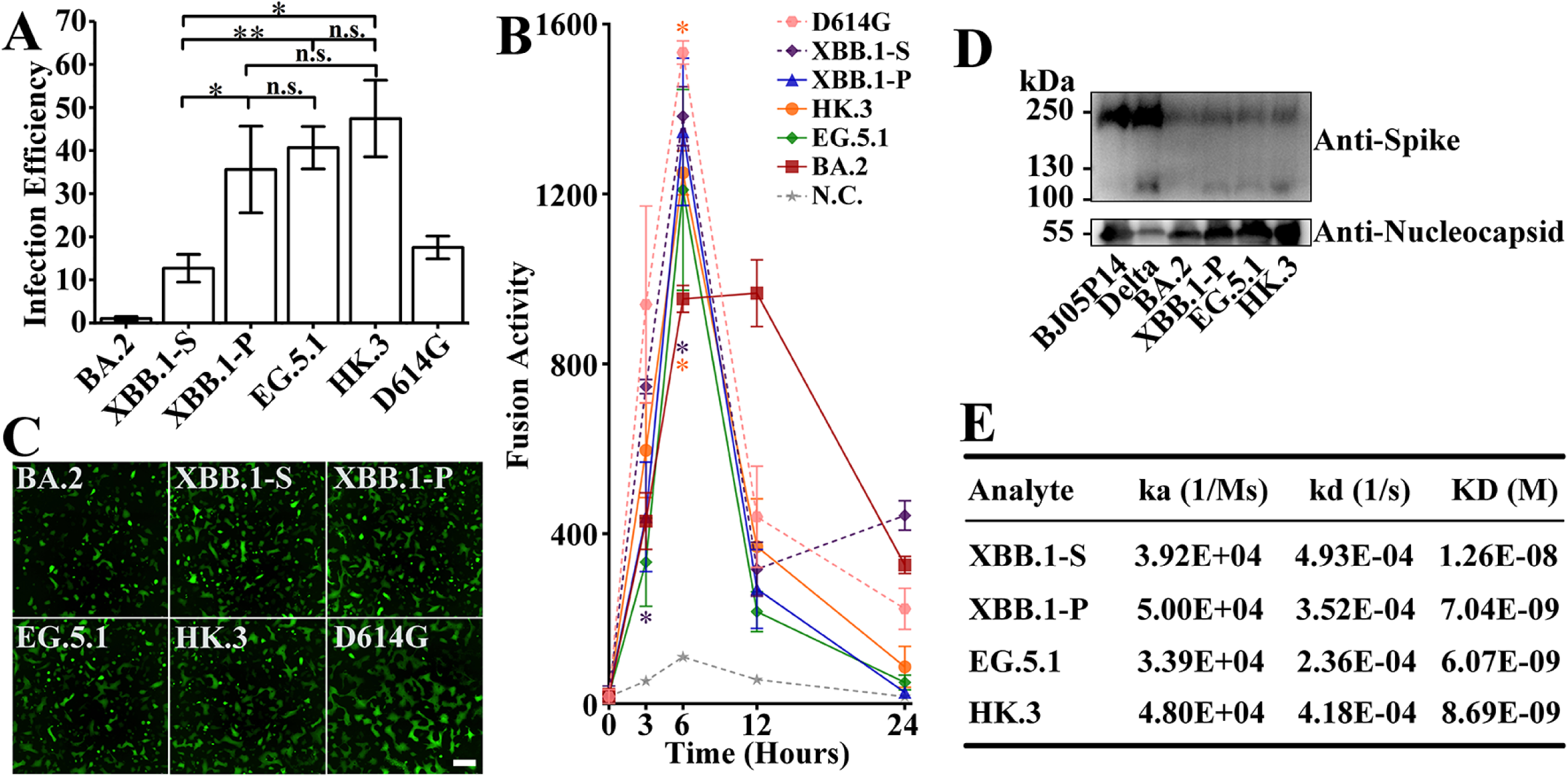
Characteristics of the spikes of XBB.1 subvariants. A. Spike-mediated infection determined by pseudovirus assays. XBB.1-F486S (referred to as XBB.1-S) of the XBB.1 lineage and XBB.1-F486P (referred to as XBB.1-P) of the XBB.1.9 lineage were included. B. Spike-mediated cell‒cell fusion based on luciferase activity. BA.2 (dark red), XBB.1-F486P (blue), EG.5.1 (green), and HK.3 (orange) are indicated by solid lines; D614G (pink), XBB.1 (purple) and a negative control (N.C. in gray) are indicated by dotted lines. The significance of the differences between XBB variants and D614G or BA.2 is indicated in colors corresponding to the individual XBB variants, which are located above or below the lines by the asterisks, respectively. C. Spike-mediated syncytia formation (scale bar: 400 μm). D. Spike processing in authentic SARS-CoV-2 virions, including BJ05P14, Delta, BA.2, XBB.1.9-F486P, EG.5.1, and HK.3 virions. E. Comparison of the binding affinities of the XBB.1 spikes to hACE2.

The activation of spike protein processing at the S1/S2 polybasic cleavage site on the virion is positively correlated with increased infection and fusogenicity (9, 11). To directly investigate the effect of the L455F/F456L double mutation on the processing of the XBB.1 spike protein, we collected authentic SARS-CoV-2 virions and performed an immunoblot assay; the results revealed ∼250 kDa bands corresponding to the full-length spike protein and ∼120 kDa bands corresponding to the S2 subunit (Fig. 2D). HK.3 with the L455F/F456L double mutations showed stronger cleavage than XBB.1-F486P (of XBB.1.9.1) and EG.5.1, and this effect was probably associated with efficient replication in the cell lines. Our results, consistent with those of a previous study (18), indicated that the Delta spike protein was strongly cleaved. Virions of another SARS-CoV-2 variant isolated in our laboratory (data not published, named BJ05P14) containing a deficient furin cleavage site in the spike protein barely exhibited spike cleavage.

Spikes of epidemic Omicron lineages have demonstrated higher binding affinities than those of previous variants (19). We purified the ectodomains of XBB.1 spikes (Fig. A2) and measured their binding affinities to hACE2 via surface plasmon resonance (SPR). XBB.1 spike proteins with F486P, including EG.5.1 and HK.3, demonstrated lower dissociation rates than XBB.1 (with F486S) and 1.5 to 2 folds higher binding affinity to hACE2 than XBB.1 (Figs. 2E and A3). Despite the slightly higher binding affinity to hACE2 of EG.5.1, the three proteins with F486P exhibited similar binding affinities for hACE2.

### Virological characteristics of XBB.1.9 subvariants in vivo

We evaluated the pathogenicity of XBB.1.9.1, EG.5.1 and HK.3 in common wild-type Syrian hamster models. All the infected hamsters survived after challenge. Although the BA.2- and EG.5.1-infected hamsters exhibited significantly lower weights than the mock-infected hamsters, most of the animals infected with the four Omicron isolates gained weight over the seven-day experiment (Fig. 3A). Although the XBB.1.9.1 and EG.5.1 groups exhibited greater viral loads in the nasal lavages than did the BA.2 group at 2 days post-infection (DPI), viral loads of all the XBB groups decreased very quickly compared to those in the BA.2 group (Fig. 3B). Thus, both XBB.1.9.1 and EG.5.1 infection led to lower viral loads in nasal lavages than that of BA.2. No significant differences in viral loads or viral titers were detected in the lungs or turbinates of the four groups at 3 DPI (Fig. 3C and D). According to the nasal lavage and turbinate results, all four Omicron infections resulted in large intragroup differences in viral titers in the lower airways, which is highly consistent with previous results from other groups and our group (20, 21).

**Fig. 3.**
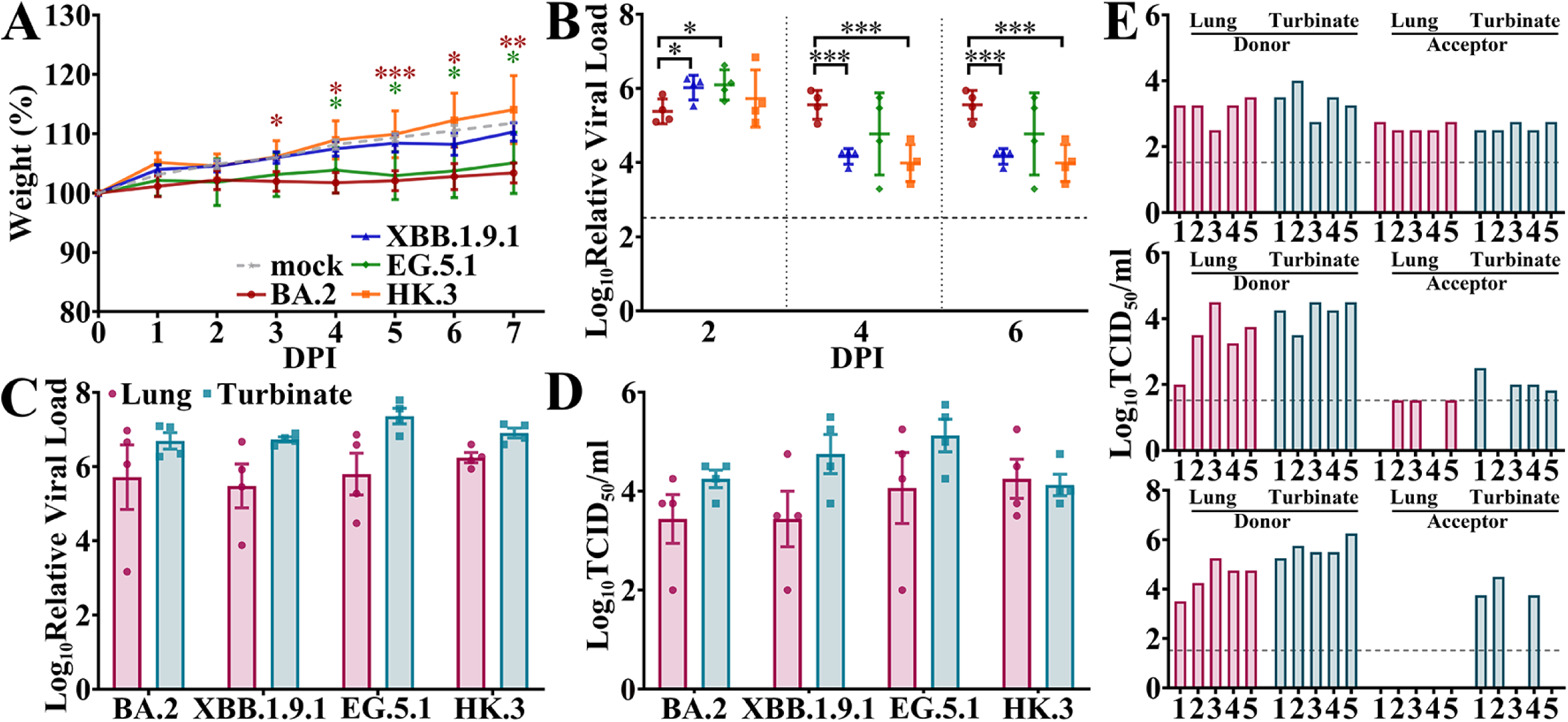
In vivo virological characteristics of the XBB.1.9 subvariants in wild-type hamsters. Hamsters were intranasally inoculated with BA.2, XBB.1.9.1, EG.5.1 or HK.3, and the corresponding results (A and B) are shown in red, blue, green and orange, respectively (as shown in A). A-B. Four hamsters per group were used to measure body weights (A) and relative viral loads in nasal lavages (B). Relative viral loads (C) and viral titers (D) in the lungs (dark red) or turbinates (cray) of the infected hamsters at 3 DPI. E. Airborne transmission of XBB.1.9.1 (upper), EG.5.1 (middle) and HK.3 (lower) in hamsters. The viral titers of the lungs and turbinates of the inoculated donors and acceptors are indicated in dark red and cray, respectively.

Given the increasing prevalence of XBB.1.9 and its sublineages, we evaluated the airborne transmission of XBB.1.9.1, EG.5.1 and HK.3 in hamsters (Fig. 3E). High transmission of XBB.1.9.1 was observed in hamsters (5/5), with infectious virus detected in both nasal turbinates and lungs. Interestingly, although EG.5.1 also exhibited high transmission in hamsters (5/5), infectious viruses were inconsistently detected in the upper and lower airways of the exposed acceptors. Specifically, lung infection of 4/5 exposed hamsters (#1, #3, #4 and #5) and turbinate infection of 3/5 hamsters (#2, #3 and #5) were demonstrated. Near-detection-limit viral titers were merely demonstrated from two acceptors (with lung of #2 and turbinate of #5). For HK.3, infectious viruses with higher titers were detected exclusively in the nasal turbinates in three of five exposed acceptors. Compared to our previous results indicating poor transmissibility of BA.2 in hamsters (1/5, exclusively detected in turbinate) (21), the transmissibility of XBB.1.9 subvariants was substantially largely enhanced, although the transmission of XBB.1.9.2 in hamsters was slightly greater than that of the other two variants.

Given the relatively efficient transmission of HK.3 and EG.5 in humans (7) and the relatively lower transmissibility in hamsters than that of XBB.1.9.1, we compared the in vivo fitness of XBB.1.9.1 and EG.5.1/HK.3 in hamsters. For animals inoculated with a 1:1 or 1:3 ratio of XBB.1.9.1:EG.5, XBB.1.9.1 showed strong fitness in the lung and turbinate samples, except for the lung of hamster 1 (Fig. 4A). Notably, with an inoculum with a 1:3 ratio of XBB.1.9.1:EG.5, the RNA proportion of XBB.1.9.1 largely increased from 54.5% to more than 70% after competitive replication in hamsters. Similarly, XBB.1.9.1 also outcompeted HK.3 in the lungs and turbinates of hamsters, except in the lungs of hamster 15 (Fig. 4B). Taken together, these results suggest that XBB.1.9.1 may have greater replicative fitness than EG.5.1 and HK.3 in hamsters, especially in the upper respiratory tract. Thus, the host-associated factor is considered one of the key points affecting XBB.1.9 transmissibility, of which the difference between human and hamster receptors may contribute. We evaluated the spike infectivity tropism of golden hamster ACE2 (ghACE2) compared to that of hACE2, however, the spikes of multiple XBB.1.9 and even BA.2 demonstrated similar tropism (Fig. A4). Therefore the differences of ACE2 receptors seem not a potential determinate of XBB transmission bias, at least revealed in hamster models.

**Fig. 4.**
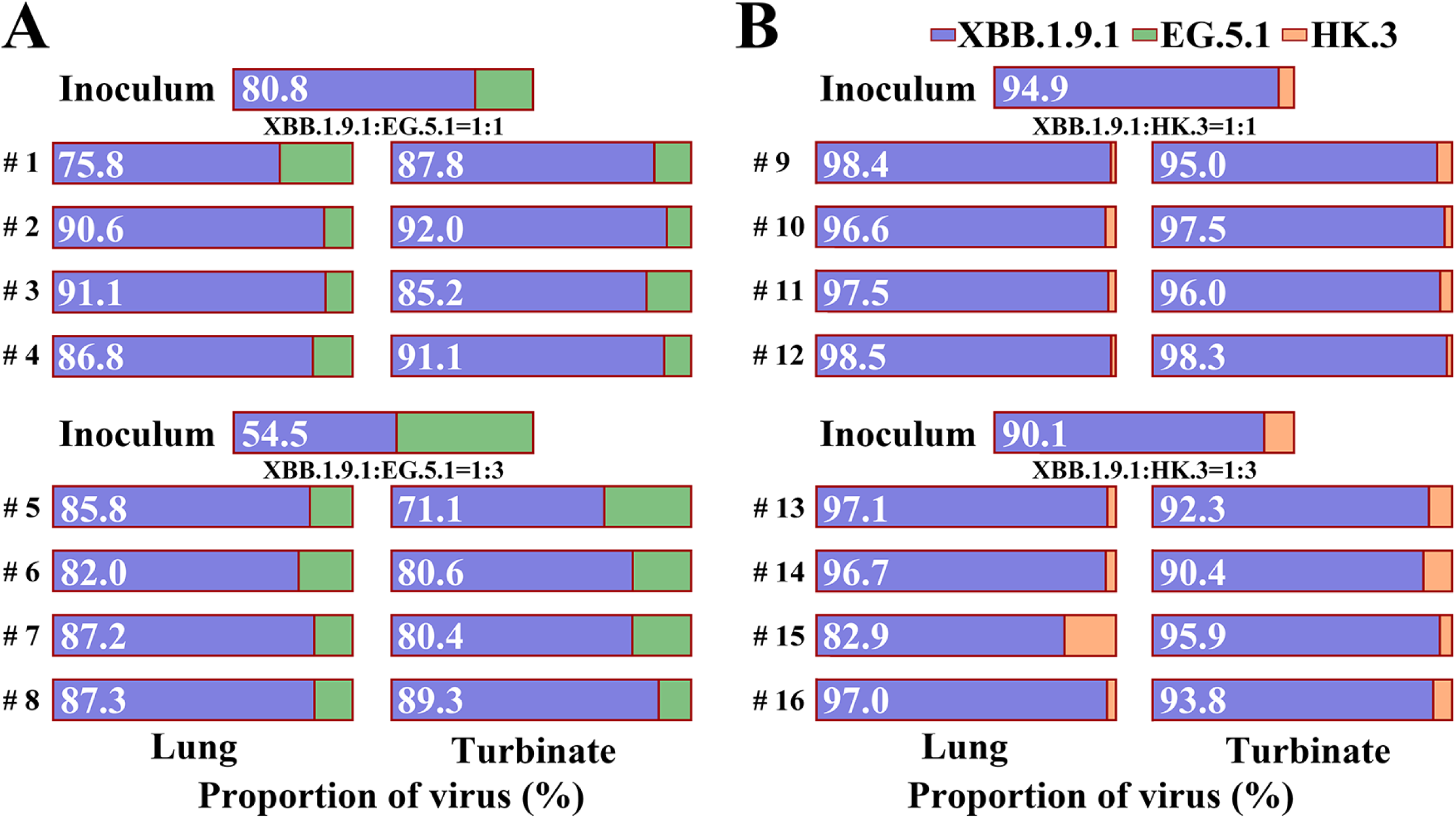
Competitive fitness of XBB.1.9.1 and EG.5/HK.3 in wild-type hamsters. XBB.1.9.1 and EG.5 (A) or HK.3 (B) at viral titer ratios of 1:1 (upper) or 1:3 (lower) were coinoculated into hamsters, and the RNA proportions of XBB.1.9.1 (blue) and EG.5.1 (green) or HK.3 (orange) in the inoculum (top), lung (left) and turbinate (right) were determined.

We then investigated the replication and pathogenicity of the XBB.1.9 subvariants by using previously reported Omicron-lethal H11-K18-hACE2 transgenic hamster models (21, 22). Similar to the results of BA.2 and BA.5.2.48 infection (21), all three XBB.1.9-infected hACE2 hamster groups exhibited high mortality (Fig. 5A). Specifically, infection with both EG.5.1 and HK.3 induced 100% lethality as early as 5 DPI, while infection with XBB.1.9.1 induced 75% lethality. BA.2 infection led to 75% and 100% lethality by 6 and 7 DPI, respectively. Compared to mock infection, XBB.1.9, EG.5 and HK.3 infection induced significant body weight changes as early as 2 DPI (Fig. 5B), which occurred earlier than that of BA.2 (at 3 DPI). Weight loss was significantly induced exclusively by EG.5.1 and HK.3 infection, which was revealed just before lethality at 4 DPI. Both the mortality and body weight changes indicated that HK.3 and EG.5.1 demonstrated greater pathogenicity than did the parental strains XBB.1.9.1 and BA.2. The XBB.1.9.1 group exhibited lower viral loads in nasal lavages than did the BA.2 group at 4 DPI, but no significant differences were found among the XBB.1.9.1, EG.5 and HK.3 groups (Fig. 5C). To evaluate the pathological features and determine the infected areas in the lungs, we performed H&E staining and anti-nucleocapsid immunohistochemistry (IHC) of the hamster lungs. We observed universal alveolar wall thickening, infiltration of inflammatory cells in alveolar spaces and bronchial mucosal epithelial cells in all groups, and the viruses caused similar infections in multiple areas of the lungs, from the epithelium of the terminal bronchiole to the alveoli. Moreover, despite the comparable pathology scores of the different groups (Fig. 5D), infection with BA.2 and XBB.1.9.1 led to greater inflammatory infiltration (see pathology scores in Fig. A5; BA.2: 2.50 and XBB.1.9.1: 2.33; compared to HK.3: 1.75 and EG.5: 1.67), while EG.5.1 and HK.3 infection tended to cause a large amount of nucleocapsid-positive exfoliation of epithelial cells in terminal bronchioles, which formed widespread obstructions in the airways (HK.3: 2.00 and EG.5: 2.33 compared to BA.2: 1.25 and XBB.1.9.1: 1.00). Interestingly, XBB.1.9.1, EG.5 and HK.3, but not BA.2, were found to infect vascular endothelial cells near alveoli (Fig. A6), producing more potential threats to more organs via the circulatory system.

**Fig. 5.**
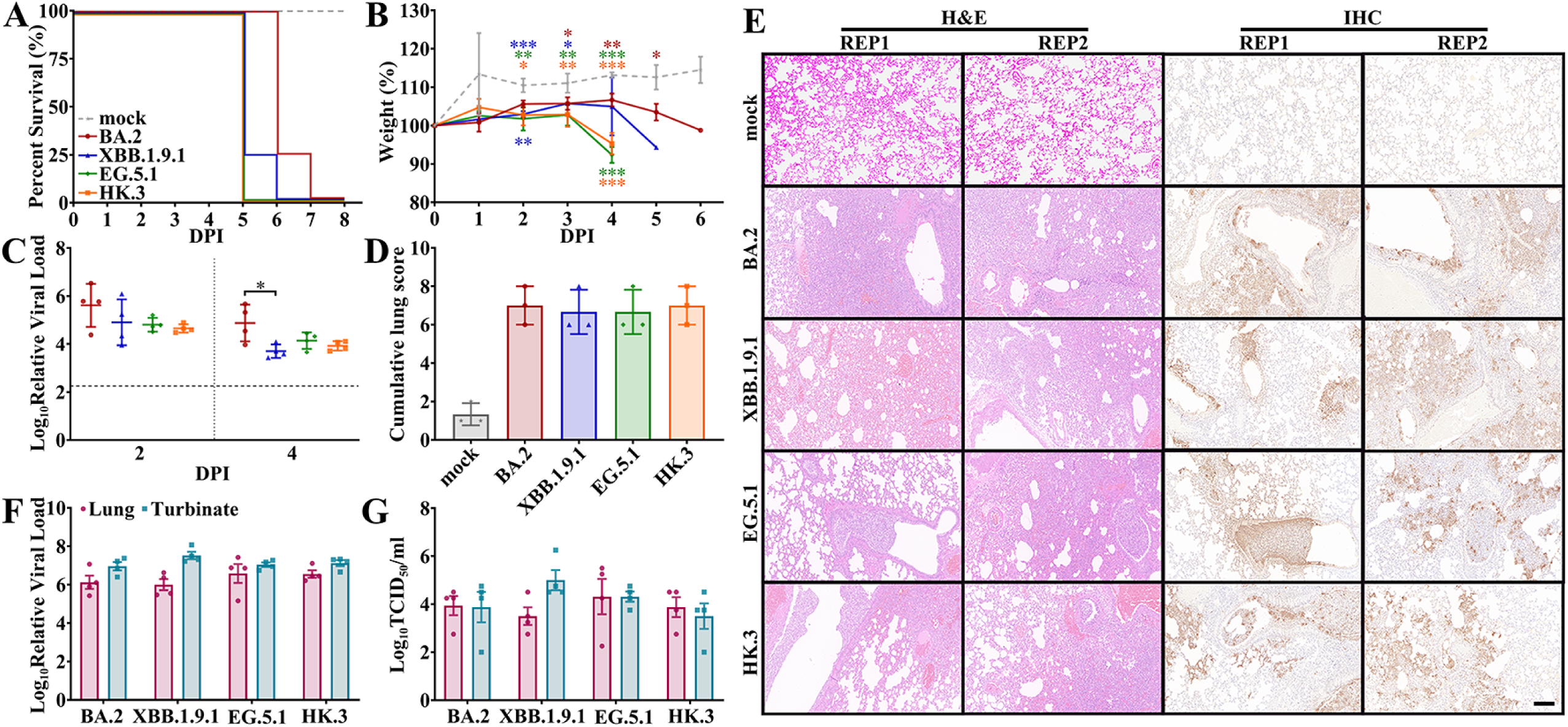
In vivo virological characteristics of XBB.1.9 subvariants in K18-hACE2 hamsters. K18-hACE2 hamsters were intranasally inoculated with BA.2, XBB.1.9.1, EG.5.1 or HK.3. Four hamsters per group were used to measure the various parameters (A, B and C). Four hamsters per group were euthanized at 3 DPI and used for data collection (D, E, F and G). The data (in A, B, C, and D) of the mock, BA.2, XBB.1.9.1, EG.5.1 and HK.3 groups are shown in gray, red, blue, green and orange, respectively (as shown in A). A. Percentage survival of the infected hamsters. B. Body weights of the infected hamsters. Significant differences between the mock group and each infected group are revealed above the lines using asterisks in the colors corresponding to the respective infected group. C. Relative viral RNA loads in the nasal lavages of hamsters. The viral load baseline is indicated by dotted gray lines. D/E. Cumulative lung scores (D) and H&E staining and IHC images (E) of the lungs of the infected hamsters. The lungs of two infected individuals in each group, namely, repetition 1 (REP1) and repetition 2 (REP2), are shown. The scale bar represents 100 μm. F/G. Relative viral RNA loads (F) and viral titers (G) in the lungs (dark red) or turbinates (cray) of the infected hamsters.

## Discussion

Omicron BA.1 and BA.2 are less likely to induce pneumonia and other severe symptoms in COVID-19 patients (23, 24), which is consistent with their attenuated pathogenicity compared with that of the previous VOCs revealed in hamster and K18-hACE2 mouse models; similarly, BA.5 also failed to significantly improve pathogenicity compared to BA.2 (20, 25, 26). However, the attenuation of pathogenicity by Omicron is not continuous. We previously reported that a further sublineage of BA.5, BA.5.2.48, demonstrated greater pathogenicity than BA.2 in H11-K18-hACE2 rodents (21). In this study, H11-K18-hACE2 hamsters infected with sublineage XBB.1.9 exhibited earlier mortality and significantly greater weight loss than did the hamsters infected with BA.2. Similarly, the spike F486P mutation led to improved pathogenicity in XBB.1 (27), which was probably attributed to its higher infectivity and hACE2 binding affinity of the spike protein reported by us and others (28). However, the infectivity, fusogenicity and hACE2 binding affinity of HK.3, EG.5.1 and XBB.1.9.1 were highly comparable (Fig. 2); thus, the reasons for the greater pathogenicity of HK.3 and EG.5.1 than of XBB.1.9.1 were not determined. In contrast to the poor airborne transmission of BA.1 and BA.2 reported by other groups and our group (21, 29), all the assessed XBB.1.9.1, EG.5.1 and HK.3 variants demonstrated efficient transmissibility in hamster models (Fig. 3E), which is probably a key factor contributing to the prevalence. Although XBB.1.9.1 exhibited stronger fitness in the upper airways, which explained to some extent its slightly greater transmissibility than that of EG.5.1 and HK.3, further evaluation on other viral and host factors (except for ACE2) that contribute to the difference in airborne transmission tropism of XBB.1.9 subvariants are also required.

## Materials and Methods

### Plasmids, PCR and qRT‒PCR

The sequences encoding the D614G, BA.2, XBB.1-S (486S), XBB.1-P (486P), EG.5.1 and HK.3 spike proteins lacking the C-terminal 19 amino acids (spike-D19) were synthesized and subcloned into a pcDNA3 plasmid (pcDNA3-spike). Codon-optimized XBB.1.9-F486P, EG.5.1 and HK.3 spike ectodomains with a furin cleavage site mutation (682RRAR685 mutated to 682GSAS685), a “HexaPro” modification and a 6×His-StrepⅡ tag were synthesized and subcloned into a pcDNA3 plasmid (pcDNA3-spike-HexaPro). The dual split protein (DSP) plasmids pDSP1-7 and pDSP8-11, which encode the split Renilla luciferase and GFP genes, were previously described (21). Viral RNA extraction was performed using a QIAamp Viral RNA Mini Kit (Qiagen). First-strand cDNA synthesis was used for reverse transcription (RT), RT‒PCR was used to amplify specific spike fragments, and qRT‒PCR was used to determine the relative genome copy number (or viral load) of SARS‒CoV‒2, as previously described (30). To distinguish replicating viral genomes in infected samples from those in aerosol-contaminated samples, we used the average viral load in samples collected from uninfected animals (mock groups) as a detection baseline in qRT‒PCR results.

### Cells, transfection, protein purification and SPR

Vero E6 cells, HeLa^hACE2+^ cells, Calu-3 cells, HEK-293T cells and HEK-293F cells were cultured and transfected using either Lipofectamine 3000 (Thermo) or polyethylenimine (Polysciences) as previously described (30, 31). HEK-293F cells were transfected with pcDNA3-spike-HexaPro for five days. Spike proteins in culture medium were purified with gravity flow columns using Strep-Tactin XT (IBA Life Sciences), concentrated to 0.5 mg/ml with ultrafiltration tubes (Millipore) and stored at -80 ℃ before use. The purified proteins were analyzed on 6% SDS‒ PAGE gels with Coomassie blue staining. The SPR experiments were performed using a Biacore T200 (Cytiva). All assays were performed with 1×HEPES running buffer (10 mM HEPES, 150 mM NaCl, 3 mM EDTA, and 0.005% Tween-20, pH 7.4) at 25℃. For determination of the binding kinetics between the spike protein and hACE2 proteins, a Protein A sensor chip (Cytiva) was used. The hACE2 protein with an Fc tag (Acro Biosystems) was immobilized onto the sample flow cell of the sensor chip. The reference flow cell was left blank. Each spike protein was injected over the two flow cells at a range of eight concentrations prepared by serial twofold dilutions (from 3.906 nM to 500 nM) at a flow rate of 30 μl/min using a single-cycle kinetics program. All the data were fitted to a 1:1 binding model using Biacore T200 Evaluation Software 3.1.

### Virus

The four SARS-CoV-2 Omicron XBB.1.9 subvariants used in this study were an XBB.1.9.1 isolate (hCoV-19/Jilin/JSY-CC8/2023, GISAID Accession No. EPI_ISL_18908494), which contains the extra mutations A10323G (NSP5_K90R), C11750T (NSP6_L260F), G25352T (Spike_V1264L) and G26634T (M_A38S); an EG.5.1 isolate (hCoV-19/Jilin/JSY-CC9/2023, GISAID Accession No. EPI_ISL_18908495), which contains the extra mutations T9037A (NSP4_D161E), T18285G (NSP14_H82Q) and G23587C (Spike_Q675H); and two HK.3 isolates, referred to as HK.3-CC11 (hCoV-19/Jilin/JSY-CC11/2023, GISAID Accession No. EPI_ISL_18908496) containing the extra mutation GTT26284-26286del (E_V14del), and HK.3-CC12 (hCoV-19/Jilin/JSY-CC12/2023, GISAID Accession No. EPI_ISL_18908497) containing the extra mutations G11083T (NSP6_L37F), C24912T (Spike_T1117I) and GTT26284-26286del (E_V14del). A BA.2 isolate (hCoV-19/Jilin/JSY-CC5/2022, GISAID Accession No. EPI_ISL_18435548) was also used. SARS-CoV-2 was passaged in Vero E6 cells, and the viral titers were measured using TCID_50_ assays as previously described (21) with a baseline of 33 TCID_50_/ml.

### Pseudovirus production and infection

The vesicular stomatitis virus (VSV) pseudotyped virus deficient in the G gene (ΔG) bearing a firefly luciferase reporter gene and the VSV-G glycoprotein for infection (VSV-Luciferase-ΔG*G, Brain Case) was passaged in HEK-293T cells transfected with the pMD2.G plasmid. For production of spike-glycoprotein-bearing pseudovirus (VSV-Luciferase-ΔG*Spike), HEK-293T cells were infected with VSV-Luciferase-ΔG*G at a multiplicity of infection (MOI) of 0.1 after transfection with pcDNA3-spike plasmids (or pcDNA3 as a negative control) and washed twice with DMEM after 2 hours. Media containing pseudoviruses at 36 HPI were centrifuged at 500 ×g for 10 min, after which the supernatant was stored at -80°C. For measurement of spike-mediated infection, 2×10^4^ HeLa^hACE2+^ cells (in 96-well culture plates) were mixed with 100 μl of pseudovirus, washed once with DMEM after 24 hours and mixed with 100 μl of luciferase substrate (PerkinElmer). Luminescence was detected using an Infinite 200 Pro plate reader (Tecan) as spike-mediated infectivity.

### Cell‒cell fusion

Spike-mediated cell‒cell fusion was determined based on the split proteins DSP8-11 and DSP1-7, which were expressed in effector and target cells, respectively. For preparation of effector cells, HEK293T cells in 12-well plates were cotransfected with 1 μg of pcDNA3-spike (or pcDNA3 as a negative control) and 1 μg of pDSP8-11. For preparation of target cells, HeLa^hACE2+^ cells in 6-well plates were transfected with pDSP1-7 (2.5 μg). A total of 1.3×10^5^ effector cells were transferred to 96-well plates at 24 hours post-transfection. After another 24 hours, the target cells were incubated with 1:250 diluted EnduRen live cell substrate (Promega). After detachment, 100 μl of 2.6×10^5^ targeted cells were added to wells containing effector cells. Luminescence was measured at the indicated time points using an Infinite 200 Pro plate reader (Tecan), and fusion activity was assessed. Cell images were taken using an APX100 microscope (Olympus).

### Animal experiments

Eight-week-old male wild-type Syrian hamsters (Charles River) or 12-week-old H11-K18-hACE2 Syrian hamsters (State Key Laboratory of Reproductive Medicine and Offspring Health, China) were intranasally inoculated with the indicated viruses at a dose of 1000 TCID_50_ (in 100 μl). Each infected group included four animals in the body weight subgroup and four animals in the euthanized subgroup for tissue sample collection.

For coinfection studies, XBB.1.9.1 was mixed with EG.5.1 or HK.3 at viral titer ratios of 1:1 or 1:3, and the virus mixture (a total of 1000 TCID_50_ in 100 µl) was inoculated into six-week-old male Syrian hamsters (Charles River). Tissue samples were collected at 3 DPI. Spike fragments flanking the shared mutations between XBB.1.9.1 and EG.5.1 or HK.3, Q52H or F456L were amplified using RT‒PCR, and the ratios were determined by geometric means of the peak height ratios in the Sanger sequencing results. The primers used are listed in Table A1.

For the airborne transmission study between hamsters, five six-week-old male hamsters were intranasally inoculated with each of the indicated viruses at a dose of 1000 TCID_50_ (in 100 μl). After 24 hours, each infected donor hamster was cohoused with one corresponding acceptor hamster in isolation devices. The propagation cages containing the donor and acceptor were separated by 3 cm to prevent direct contact. The inoculated hamsters were placed in front of the isolator unit, which provides unidirectional airflow. Tissue samples were collected 3 days after infection for the donor hamsters or 3 days after the initial cohousing for the exposed acceptors.

For all animals, intranasal inoculation and euthanasia were performed under isoflurane anesthesia.

### Histopathology

The lungs of the animals were fixed in 4% paraformaldehyde in phosphate-buffered saline (PBS) and processed for paraffin embedding. The paraffin blocks were sliced into 3-µm thick sections and mounted on silane-coated glass slides, which were subsequently subjected to H&E staining for histopathological examination. Panoramic images of the digital slides were taken using a Pannoramic MIDI (3DHISTECH). Pathological features, including (i) bronchiolitis or exfoliation of epithelial cells in terminal bronchioles; (ii) hemorrhage; (iii) alveolar damage, including alveolar thickening or disappearance; and (iv) inflammatory infiltration, were scored using a four-tiered system as 0 (negative), 1 (weak), 2 (moderate), or 3 (severe).

### Immunoblotting, IHC and immunofluorescence

Virions of SARS-CoV-2 in clear supernatant (10 ml) were inactivated by adding 0.1% paraformaldehyde (final concentration) for 12 hours at 4°C and then concentrated by ultracentrifugation at 90000 ×g for 2 hours using a P80A rotor (Hettich). The pellets were resuspended in 30 μl of RIPA buffer. The samples were separated via 4-12% SDS‒PAGE gels and transferred to PVDF membranes. After the membranes were blocked with 5% milk, they were blotted with primary antibodies, incubated with horseradish peroxidase-conjugated secondary antibodies and visualized with a chemiluminescent reagent as previously described^35^. For IHC, tissue sections were processed for IHC with a primary antibody and horseradish peroxidase-conjugated secondary antibody and finally stained with a DAB substrate kit (Solarbio). Panoramic images of the digital slides were taken using an SQS-40R slide scan system (Shengqiang Technology). Immunofluorescence was performed as previously described^35^. The antibodies used are listed in Table A2.

### Statistical analysis

All the data are presented as the means ± SDs. Comparisons were performed using Student’s tests. Significance was defined as P<0.05 (*), P<0.01 (**) or P<0.001 (***) and was indicated either in the figures or mentioned separately in the text. The number of repeats is specified in individual panels using discrete points. All presented data are biological replicates.

### Ethics and biosecurity

All animal experiments were approved by the Animal Care and Use Committee of the Changchun Veterinary Research Institute (approval number: AMMS-11-2023-032). All experiments involving infectious SARS-CoV-2 were performed at the Animal Biosafety Level 3 Laboratories of the Changchun Veterinary Research Institute, Chinese Academy of Agricultural Sciences.

## Acknowledgments

Data sharing statement

The original data that support the findings of this study are available from the corresponding author upon reasonable request.

## Fundings

This work is supported by the National Key Research and Development Program of China (2021YFC2302405 to X.W., 2023YFC0871100 to Y.G., 2021YFF0702500 to J.L. and 2023YFC2605500 to Q.C.). This work is also supported by the Natural Science Foundation of Jiangsu Province (BE2019730) and Medical Research Project of Jiangsu Provincial Health Commission (Z2023003) to W.Z.

## Author Contributions

Q.J., R.L., W.W., H.Z. T.W., and H.X. performed all the virus-related experiments. W.Z., Q.C., Y.G., and Y.L. generated the H11-K18-hACE2 hamsters. X.W. designed the study. Q.J and X.W. wrote the first draft. Y.G., J.L., F.Y., and X.X. revised the first draft and approved the submitted version.

## Declaration of interests

The authors declare no competing interests.

## Appendixes

Figs. A1 to A6 and Tables A1 to A2 are obtained from Appendixes.

## References

1. Gangavarapu K, Latif AA, Mullen JL, Alkuzweny M, Hufbauer E, Tsueng G, Haag E, Zeller M, Aceves CM, Zaiets K, Cano M, Zhou X, Qian Z, Sattler R, Matteson NL, Levy JI, Lee RTC, Freitas L, Maurer-Stroh S, Suchard MA, Wu C, Su AI, Andersen KG, Hughes LD. 2023. Outbreak.info genomic reports: scalable and dynamic surveillance of SARS-CoV-2 variants and mutations. Nature Methods 20:512–522.

2. Wang Q, Iketani S, Li Z, Liu L, Guo Y, Huang Y, Bowen AD, Liu M, Wang M, Yu J, Valdez R, Lauring AS, Sheng Z, Wang HH, Gordon A, Liu L, Ho DD. 2023. Alarming antibody evasion properties of rising SARS-CoV-2 BQ and XBB subvariants. Cell 186:279–286.e8.

3. Uraki R, Ito M, Furusawa Y, Yamayoshi S, Iwatsuki-Horimoto K, Adachi E, Saito M, Koga M, Tsutsumi T, Yamamoto S, Otani A, Kiso M, Sakai-Tagawa Y, Ueki H, Yotsuyanagi H, Imai M, Kawaoka Y. 2023. Humoral immune evasion of the omicron subvariants BQ.1.1 and XBB. The Lancet Infectious Diseases 23:30–32.

4. Qu P, Faraone JN, Evans JP, Zheng Y-M, Carlin C, Anghelina M, Stevens P, Fernandez S, Jones D, Panchal AR, Saif LJ, Oltz EM, Zhang B, Zhou T, Xu K, Gumina RJ, Liu S-L. 2023. Enhanced evasion of neutralizing antibody response by Omicron XBB.1.5, CH.1.1, and CA.3.1 variants. Cell Reports 42:112443.

5. Kurhade C, Zou J, Xia H, Liu M, Chang HC, Ren P, Xie X, Shi PY. 2023. Low neutralization of SARS-CoV-2 Omicron BA.2.75.2, BQ.1.1 and XBB.1 by parental mRNA vaccine or a BA.5 bivalent booster. Nature Medicine 29:344–347.

6. Faraone JN, Qu P, Goodarzi N, Zheng Y-M, Carlin C, Saif LJ, Oltz EM, Xu K, Jones D, Gumina RJ, Liu S-L. 2023. Immune evasion and membrane fusion of SARS-CoV-2 XBB subvariants EG.5.1 and XBB.2.3. Emerging Microbes & Infections 12.

7. Kosugi Y, Plianchaisuk A, Putri O, Uriu K, Kaku Y, Hinay AA, Jr., Chen L, Kuramochi J, Sadamasu K, Yoshimura K, Asakura H, Nagashima M, Ito J, Misawa N, Guo Z, Tolentino JEM, Fujita S, Pan L, Suganami M, Chiba M, Yoshimura R, Yasuda K, Iida K, Ohsumi N, Strange AP, Tanaka S, Fukuhara T, Tamura T, Suzuki R, Suzuki S, Ito H, Matsuno K, Sawa H, Nao N, Tanaka S, Tsuda M, Wang L, Oda Y, Ferdous Z, Shishido K, Nakagawa S, Shirakawa K, Takaori-Kondo A, Nagata K, Nomura R, Horisawa Y, Tashiro Y, Kawai Y, Takayama K, Hashimoto R, et al. Characteristics of the SARS-CoV-2 omicron HK.3 variant harbouring the FLip substitution. The Lancet Microbe doi:10.1016/S2666-5247(23)00373-7.

8. Zhou B, Thao TTN, Hoffmann D, Taddeo A, Ebert N, Labroussaa F, Pohlmann A, King J, Steiner S, Kelly JN, Portmann J, Halwe NJ, Ulrich L, Trüeb BS, Fan X, Hoffmann B, Wang L, Thomann L, Lin X, Stalder H, Pozzi B, de Brot S, Jiang N, Cui D, Hossain J, Wilson MM, Keller MW, Stark TJ, Barnes JR, Dijkman R, Jores J, Benarafa C, Wentworth DE, Thiel V, Beer M. 2021. SARS-CoV-2 spike D614G change enhances replication and transmission. Nature 592:122–127.

9. Suzuki R, Yamasoba D, Kimura I, Wang L, Kishimoto M, Ito J, Morioka Y, Nao N, Nasser H, Uriu K, Kosugi Y, Tsuda M, Orba Y, Sasaki M, Shimizu R, Kawabata R, Yoshimatsu K, Asakura H, Nagashima M, Sadamasu K, Yoshimura K, Suganami M, Oide A, Chiba M, Ito H, Tamura T, Tsushima K, Kubo H, Ferdous Z, Mouri H, Iida M, Kasahara K, Tabata K, Ishizuka M, Shigeno A, Tokunaga K, Ozono S, Yoshida I, Nakagawa S, Wu J, Takahashi M, Kaneda A, Seki M, Fujiki R, Nawai BR, Suzuki Y, Kashima Y, Abe K, Imamura K, Shirakawa K, et al. 2022. Attenuated fusogenicity and pathogenicity of SARS-CoV-2 Omicron variant. Nature 603:700–705.

10. Barut GT, Halwe NJ, Taddeo A, Kelly JN, Schön J, Ebert N, Ulrich L, Devisme C, Steiner S, Trüeb BS, Hoffmann B, Veiga IB, Leborgne NGF, Moreira EA, Breithaupt A, Wylezich C, Höper D, Wernike K, Godel A, Thomann L, Flück V, Stalder H, Brügger M, Esteves BIO, Zumkehr B, Beilleau G, Kratzel A, Schmied K, Ochsenbein S, Lang RM, Wider M, Machahua C, Dorn P, Marti TM, Funke-Chambour M, Rauch A, Widera M, Ciesek S, Dijkman R, Hoffmann D, Alves MP, Benarafa C, Beer M, Thiel V. 2022. The spike gene is a major determinant for the SARS-CoV-2 Omicron-BA.1 phenotype. Nature Communications 13:5929.

11. Escalera A, Gonzalez-Reiche AS, Aslam S, Mena I, Laporte M, Pearl RL, Fossati A, Rathnasinghe R, Alshammary H, van de Guchte A, Farrugia K, Qin Y, Bouhaddou M, Kehrer T, Zuliani-Alvarez L, Meekins DA, Balaraman V, McDowell C, Richt JA, Bajic G, Sordillo EM, Dejosez M, Zwaka TP, Krogan NJ, Simon V, Albrecht RA, van Bakel H, García-Sastre A, Aydillo T. 2022. Mutations in SARS-CoV-2 variants of concern link to increased spike cleavage and virus transmission. Cell Host & Microbe 30:373–387.e7.

12. Pung R, Kong XP, Cui L, Chae S-R, Chen MIC, Lee VJ, Ho ZJM. 2023. Severity of SARS-CoV-2 Omicron XBB subvariants in Singapore. The Lancet Regional Health - Western Pacific 37:100849.

13. Valanparambil RM, Lai L, Johns MA, Davis-Gardner M, Linderman SL, McPherson TO, Chang A, Akhtar A, Gamarra ELB, Matia H, McCook-Veal AA, Switchenko J, Nasti TH, Green F, Saini M, Wieland A, Pinsky BA, Solis D, Dhodapkar MV, Carlisle J, Ramalingam S, Ahmed R, Suthar MS. 2023. BA.5 bivalent booster vaccination enhances neutralization of XBB.1.5, XBB.1.16 and XBB.1.9 variants in patients with lung cancer. npj Vaccines 8.

14. Liu Y, Liu J, Plante KS, Plante JA, Xie X, Zhang X, Ku Z, An Z, Scharton D, Schindewolf C, Widen SG, Menachery VD, Shi P-Y, Weaver SC. 2022. The N501Y spike substitution enhances SARS-CoV-2 infection and transmission. Nature 602:294–299.

15. Furusawa Y, Kiso M, Iida S, Uraki R, Hirata Y, Imai M, Suzuki T, Yamayoshi S, Kawaoka Y. 2023. In SARS-CoV-2 delta variants, Spike-P681R and D950N promote membrane fusion, Spike-P681R enhances spike cleavage, but neither substitution affects pathogenicity in hamsters. eBioMedicine 91:104561.

16. Bussani R, Schneider E, Zentilin L, Collesi C, Ali H, Braga L, Volpe MC, Colliva A, Zanconati F, Berlot G, Silvestri F, Zacchigna S, Giacca M. 2020. Persistence of viral RNA, pneumocyte syncytia and thrombosis are hallmarks of advanced COVID-19 pathology. EBioMedicine 61:103104.

17. Lin L, Li Q, Wang Y, Shi Y. 2021. Syncytia formation during SARS-CoV-2 lung infection: a disastrous unity to eliminate lymphocytes. Cell Death & Differentiation 28:2019–2021.

18. Saito A, Irie T, Suzuki R, Maemura T, Nasser H, Uriu K, Kosugi Y, Shirakawa K, Sadamasu K, Kimura I, Ito J, Wu J, Iwatsuki-Horimoto K, Ito M, Yamayoshi S, Loeber S, Tsuda M, Wang L, Ozono S, Butlertanaka EP, Tanaka YL, Shimizu R, Shimizu K, Yoshimatsu K, Kawabata R, Sakaguchi T, Tokunaga K, Yoshida I, Asakura H, Nagashima M, Kazuma Y, Nomura R, Horisawa Y, Yoshimura K, Takaori-Kondo A, Imai M, Chiba M, Furihata H, Hasebe H, Kitazato K, Kubo H, Misawa N, Morizako N, Noda K, Oide A, Suganami M, Takahashi M, Tsushima K, Yokoyama M, Yuan Y, et al. 2022. Enhanced fusogenicity and pathogenicity of SARS-CoV-2 Delta P681R mutation. Nature 602:300–306.

19. Xu Y, Wu C, Cao X, Gu C, Liu H, Jiang M, Wang X, Yuan Q, Wu K, Liu J, Wang D, He X, Wang X, Deng S-J, Xu HE, Yin W. 2022. Structural and biochemical mechanism for increased infectivity and immune evasion of Omicron BA.2 variant compared to BA.1 and their possible mouse origins. Cell Research 32:609–620.

20. Uraki R, Halfmann PJ, Iida S, Yamayoshi S, Furusawa Y, Kiso M, Ito M, Iwatsuki-Horimoto K, Mine S, Kuroda M, Maemura T, Sakai-Tagawa Y, Ueki H, Li R, Liu Y, Larson D, Fukushi S, Watanabe S, Maeda K, Pekosz A, Kandeil A, Webby RJ, Wang Z, Imai M, Suzuki T, Kawaoka Y. 2022. Characterization of SARS-CoV-2 Omicron BA.4 and BA.5 isolates in rodents. Nature 612:540–545.

21 . Wang W, Jin Q, Liu R, Zeng W, Zhu P, Li T, Wang T, Xiang H, Zhang H, Chen Q, Gao Y, Lai Y, Yan F, Xia X, Li J, Wang X, Gao Y. 2024. Virological characteristics of SARS-CoV-2 Omicron BA.5.2.48. bioRxiv doi:10.1101/2024.03.26.586802:2024.03.26.586802.

22. Dong M, Lin H, Pan M, Huang M, Liu M, Jiang R, Lai Y, Shi A, Yao B, Hu B, Shi Z, Zhang A, Gao Y, Zeng W, Jianmin L. 2024. Characterization of the SARS-CoV-2 BA.5 Variants in H11-K18-hACE2 Hamsters. bioRxiv doi:10.1101/2024.02.19.581112:2024.02.19.581112.

23. Espenhain L, Funk T, Overvad M, Edslev SM, Fonager J, Ingham AC, Rasmussen M, Madsen SL, Espersen CH, Sieber RN, Stegger M, Gunalan V, Wilkowski B, Larsen NB, Legarth R, Cohen AS, Nielsen F, Lam JUH, Lavik KE, Karakis M, Spiess K, Marving E, Nielsen C, Wiid Svarrer C, Bybjerg-Grauholm J, Olsen SS, Jensen A, Krause TG, Müller L. 2021. Epidemiological characterisation of the first 785 SARS-CoV-2 Omicron variant cases in Denmark, December 2021. Eurosurveillance 26:2101146.

24. Li X, Wu L, Qu Y, Cao M, Feng J, Huang H, Liu Y, Lu H, Liu Q, Liu Y. 2022. Clinical characteristics and vaccine effectiveness against SARS-CoV-2 Omicron subvariant BA.2 in the children. Signal Transduction and Targeted Therapy 7:203.

25. Halfmann PJ, Iida S, Iwatsuki-Horimoto K, Maemura T, Kiso M, Scheaffer SM, Darling TL, Joshi A, Loeber S, Singh G, Foster SL, Ying B, Case JB, Chong Z, Whitener B, Moliva J, Floyd K, Ujie M, Nakajima N, Ito M, Wright R, Uraki R, Warang P, Gagne M, Li R, Sakai-Tagawa Y, Liu Y, Larson D, Osorio JE, Hernandez-Ortiz JP, Henry AR, Ciuoderis K, Florek KR, Patel M, Odle A, Wong L-YR, Bateman AC, Wang Z, Edara V-V, Chong Z, Franks J, Jeevan T, Fabrizio T, DeBeauchamp J, Kercher L, Seiler P, Gonzalez-Reiche AS, Sordillo EM, Chang LA, van Bakel H, et al. 2022. SARS-CoV-2 Omicron virus causes attenuated disease in mice and hamsters. Nature 603:687–692.

26. Shuai H, Chan JF-W, Hu B, Chai Y, Yoon C, Liu H, Liu Y, Shi J, Zhu T, Hu J-C, Hu Y-f, Hou Y, Huang X, Yuen TT-T, Wang Y, Zhang J, Xia Y, Chen L-L, Cai J-P, Zhang AJ, Yuan S, Zhou J, Zhang B-Z, Huang J-D, Yuen K-Y, To KK-W, Chu H. 2023. The viral fitness and intrinsic pathogenicity of dominant SARS-CoV-2 Omicron sublineages BA.1, BA.2, and BA.5. eBioMedicine 95.

27. Tamura T, Irie T, Deguchi S, Yajima H, Tsuda M, Nasser H, Mizuma K, Plianchaisuk A, Suzuki S, Uriu K, Begum MM, Shimizu R, Jonathan M, Suzuki R, Kondo T, Ito H, Kamiyama A, Yoshimatsu K, Shofa M, Hashimoto R, Anraku Y, Kimura KT, Kita S, Sasaki J, Sasaki-Tabata K, Maenaka K, Nao N, Wang L, Oda Y, Sawa H, Kawabata R, Watanabe Y, Sakamoto A, Yasuhara N, Suzuki T, Nakajima Y, Ferdous Z, Shishido K, Mugita Y, Takahashi O, Ichihara K, Kaku Y, Misawa N, Guo Z, Hinay A, Kosugi Y, Fujita S, Tolentino JM, Chen L, Pan L, et al. 2024. Virological characteristics of the SARS-CoV-2 Omicron XBB.1.5 variant. Nature Communications 15.

28. Tamura T, Ito J, Uriu K, Zahradnik J, Kida I, Anraku Y, Nasser H, Shofa M, Oda Y, Lytras S, Nao N, Itakura Y, Deguchi S, Suzuki R, Wang L, Begum MSTM, Kita S, Yajima H, Sasaki J, Sasaki-Tabata K, Shimizu R, Tsuda M, Kosugi Y, Fujita S, Pan L, Sauter D, Yoshimatsu K, Suzuki S, Asakura H, Nagashima M, Sadamasu K, Yoshimura K, Yamamoto Y, Nagamoto T, Schreiber G, Maenaka K, Ito H, Misawa N, Kimura I, Suganami M, Chiba M, Yoshimura R, Yasuda K, Iida K, Ohsumi N, Strange AP, Takahashi O, Ichihara K, Shibatani Y, Nishiuchi T, et al. 2023. Virological characteristics of the SARS-CoV-2 XBB variant derived from recombination of two Omicron subvariants. Nature Communications 14:2800.

29. Boon ACM, Darling TL, Halfmann PJ, Franks J, Webby RJ, Barouch DH, Port JR, Munster VJ, Diamond MS, Kawaoka Y. 2022. Reduced airborne transmission of SARS-CoV-2 BA.1 Omicron virus in Syrian hamsters. PLOS Pathogens 18:e1010970.

30. Wang X, Wang W, Wang T, Wang J, Jiang Y, Wang X, Qiu Z, Feng N, Sun W, Li C, Yang S, Xia X, He H, Gao Y. 2023. SARS-CoV-2 ORF8 Protein Induces Endoplasmic Reticulum Stress-like Responses and Facilitates Virus Replication by Triggering Calnexin: an Unbiased Study. Journal of Virology 97:e00011–23.

31. Wang X, Zhao Y, Yan F, Wang T, Sun W, Feng N, Wang W, Wang H, He H, Yang S, Xia X, Gao Y. 2021. Viral and Host Transcriptomes in SARS-CoV-2-Infected Human Lung Cells. Journal of Virology 95:e00600–21.

